## Supplementary Material for "Nitric Oxide controls shoot meristem activity via regulation of DNA methylation"

[2] Current address: Biotechnology Research Institute, Chinese Academy of Agricultural Sciences, Beijing 100081, China.

[3] Current address: CureVac, 72076 Tübingen, Germany

**This PDF file includes:**

**Supplementary Methods**

**Supplementary Figures 1 to 21**

**Supplementary References**

### Supplementary Methods

**Quantitative RT-PCR.** Whole seedlings or inflorescence apices were dissected as previously described and were immediately transferred to liquid nitrogen[1]. Total RNA was performed using the RNAeasy Plant Mini Kit following the manufacturer instructions (Qiagen). cDNA was prepared using cDNA synthesis kit after DNase treatment (Thermo Fischer). Primers used for RT-qPCR were designed for amplifying products of 100 to 300 bp in length, and primer sequences are listed in Supplementary Table 7. Quantitative PCR was performed using SYBR Green kit (EurX) with the following PCR conditions: Step 1, 95°C for 5 mins; Step 2, 40 cycles of 95°C for 10 s followed by 57°C for 30 s and 72°C for 30 s; Step 3, 72°C for 10 mins; and Step 4, 20°C for 10 s. *PP2A* was used to normalize mRNA levels.

**Co-IP assays.** Each construct was introduced into *Agrobacterium* ASE cells, which were used subsequently to infiltrate leaves of Nicotiana benthamiana. For co-IP, nuclear proteins were isolated from formaldehyde-fixed leaves. Immunoprecipitation was carried out with GFP-Trap agarose beads following manufacturer instructions. GFP- or mCherry-tagged proteins in the input and immunoprecipitated were detected by immunoblotting using anti-GFP antibody or anti-mCherry antibody (abcam).

**Heatmaps.** Heatmaps were generated using the “heatmap” or “heatmap2” program in R project, version 3.1.0. The software packages were downloaded from Bioconductor (http://bioconductor.org). The values of the relative expression levels of the genes for heatmaps consisted of log2-transformed data.

**eChIP**. The eChIP experiments were performed as described[2]. Briefly, 0.1g 7-day old seedlings were crosslinked with 1% formaldehyde for 15 mins and quenched with 0.2M glycine at room temperature. A Diagenode Bioruptor UCD-200 was used for sonication and the ChromoTek RFP-Trap Magnetic antibody was used to precipitate chromatin.

**Statistical analyses.** For the multiple sample variance statistical analysis, the significance level was obtained using a one-way fixed-effects ANOVA model as described by Larson et al.[3]. Differences between groups were identified using Student's t-test, and the p-value level was set at 5%.

### Supplementary Figures and Legends

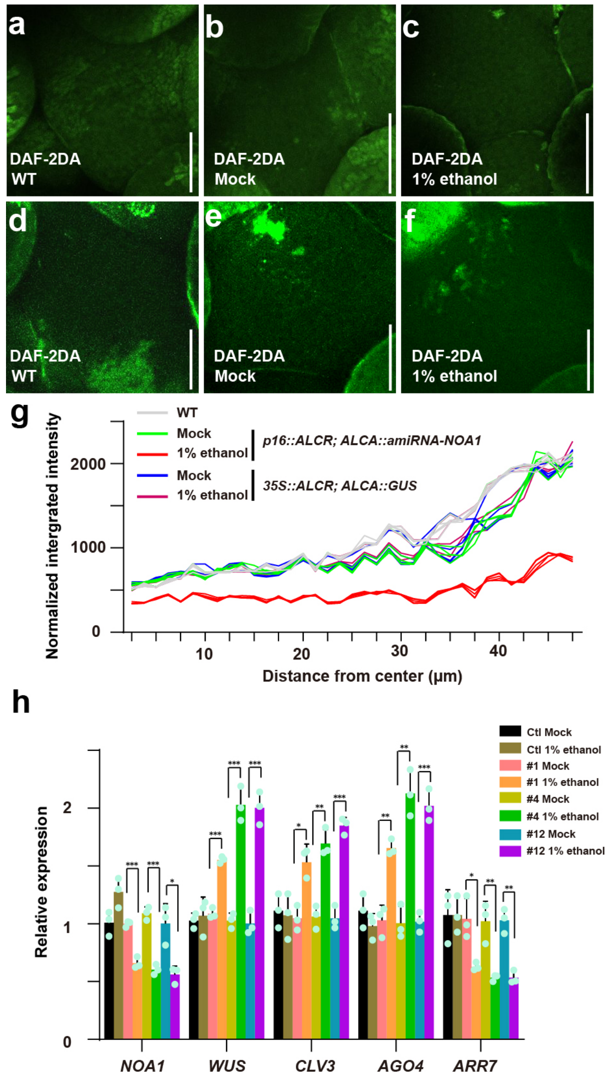


### Supplementary Fig. 1 | Transient decrease of NO levels affects stem cell regulation.

**a-c**, DAF-2DA staining in wild type (**a**) and in *p16::AlcR; pAlcA::amiRNA-NOA1* transgenic plants with (**c**) and without (**b**) 1% ethanol induction. Scale bars, 50 μm. **d-f**, DAF-2DA staining in wild type (**d**) and in the *p35S::AlcR; pAlcA::GUS* transgenic plants with (**f**) and without (**e**) 1% ethanol induction. Scale bars, 50 μm. **g**, Quantification of DAF-2DA fluorescent intensity in wild type and *p16::ALCR; pAlcA::amiRNA-NOA1* and *p35S::ALCR; pAlcA::GUS* transgenic plants with or without 1% ethanol induction. **h**, RT-qPCR quantification of *NOA1*, *WUS*, *CLV3*, *AGO4* and *ARR7* expression levels in the control (Ctl) (*p35S::AlcR;pAlcA::GUS*) and *p16::AlcR;pAlcA::amiRNA-NOA1* transgenic plants with or without 1% ethanol induction. Data are shown as mean ±s.d.; n=3 biological replicates, two-tailed Student's t-tests, *p<0.05, **p<0.01, ***p<0.001. All experiments were repeated at least two times (each time using multiple shoot apices from multiple plants), with similar results.

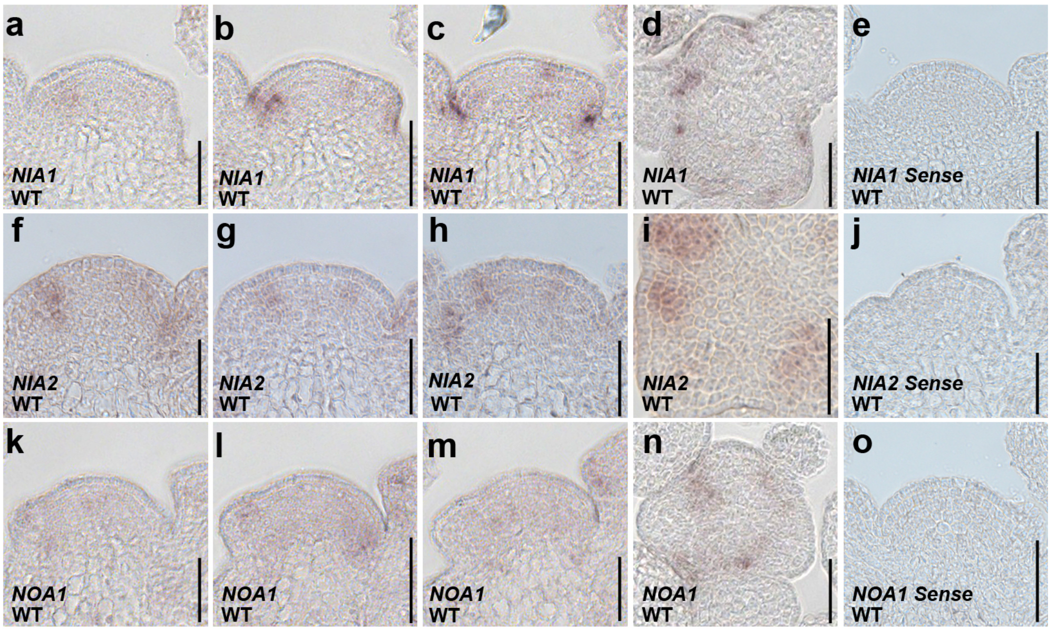


**Supplementary Fig. 2 | Expression patterns of NO biosynthesis genes in wild-type SAMs.**

**a**-**d**, *NIA1* expression patterns in the SAM at the reproductive stage (**a-c**, serial longitudinal sections; **d**, transverse section). Scale bars, 50 µm. **e**, Negative control using *NIA1* sense probe. Scale bar, 50 µm. **f-i**, *NIA2* expression patterns in the SAM at the reproductive stage (**f-h**, serial longitudinal sections; **i**, transverse section). Scale bars, 50 µm. **j**, Negative control using *NIA2* sense probe. Scale bar, 50 µm. **k-n**, *NOA1* expression patterns in the SAM at the reproductive stage (**k-m**, serial longitudinal sections; **n**, transverse section). Scale bars, 50 µm. **o**, Negative control using *NOA1* sense probe. Scale bar, 50 µm.

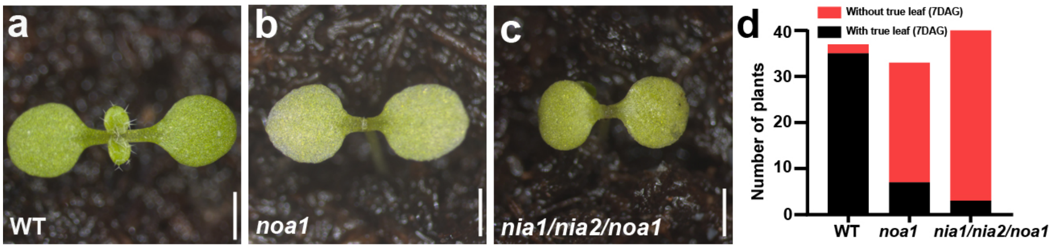


**Supplementary Fig. 3 | Nitric oxide is required for seedling development.**

**a**-**c**, Representative phenotypes of wild type (**a**), *noa1* (**b**), and *nia1/nia2/noa1* plants (**c**) seedlings seven days after germination. Scale bars, 1 mm. **d**, The percentage of wild type, *noa1* and *nia1/nia2/noa1* plants with the first pair of true leaves 7 days after germination.

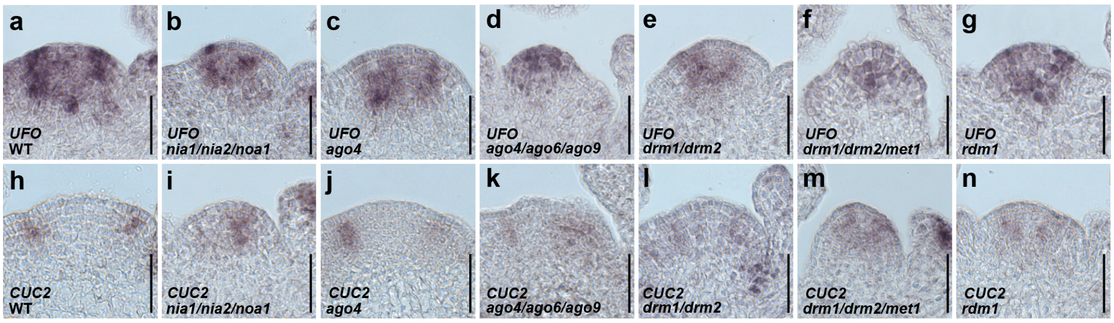


**Supplementary Fig. 4 | Expression patterns of *UFO* and *CUC2* in NO and RdDM mutants.**

**a**-**g**, *UFO* expression patterns in wild type (**a**) and in *nia1/nia2/noa1* (**b**), *ago4* (**c**), *ago4/ago6/ago9* (**d**), *drm1/drm2* (**e**), *drm1/drm2/met1* (**f**), *rdm1* mutants (**g**). Scale bars, 50 µm. **h**-**n**, *CUC2* expression patterns in wild type (**h**) and in *nia1/nia2/noa1* (**i**), *ago4* (**j**), *ago4/ago6/ago9* (**k**), *drm1/drm2* (**l**), *drm1/drm2/met1* (**m**), *rdm1* mutants (**n**). Scale bars, 50 µm.

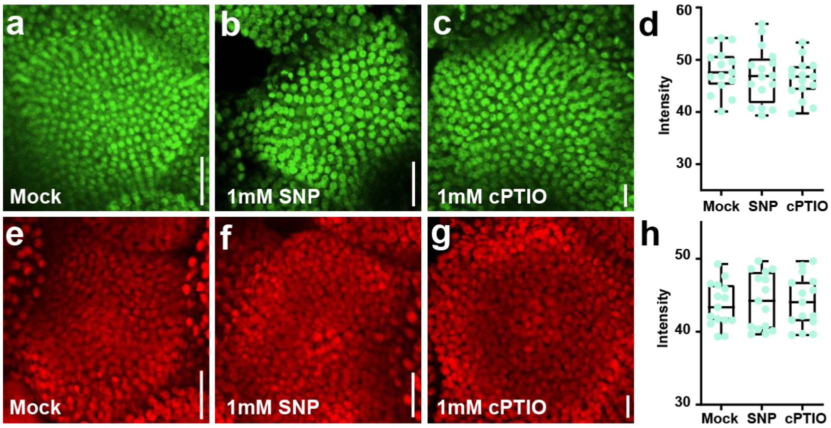


**Supplementary Fig. 5 | NO donor and scavenger treatments do not affect the activity of fluorescent proteins *in vivo*.**

**a-c**, In the *pH3.3::GFP* transgenic plants, the intensity of the GFP signals were examined using the treatments with Mock (**a**), 1mM SNP (**b**) and 1mM c-PTIO (**c**). Scale bars, 50 µm. **d**, Quantification of GFP fluorescent intensity in (a-c). n=15, ***P < 0.001, Student's t-test. **e-g**, In the *pUBQ10::mCherry* transgenic plants, the intensity of the mCherry signals were examined using the treatments with Mock (**e**), 1mM SNP (**f**) and 1mM c-PTIO (**g**). Scale bars, 50 µm. **h**, Quantification of mCherry fluorescent intensity in (e-g). n=15, ***P < 0.001, Student's t-test.

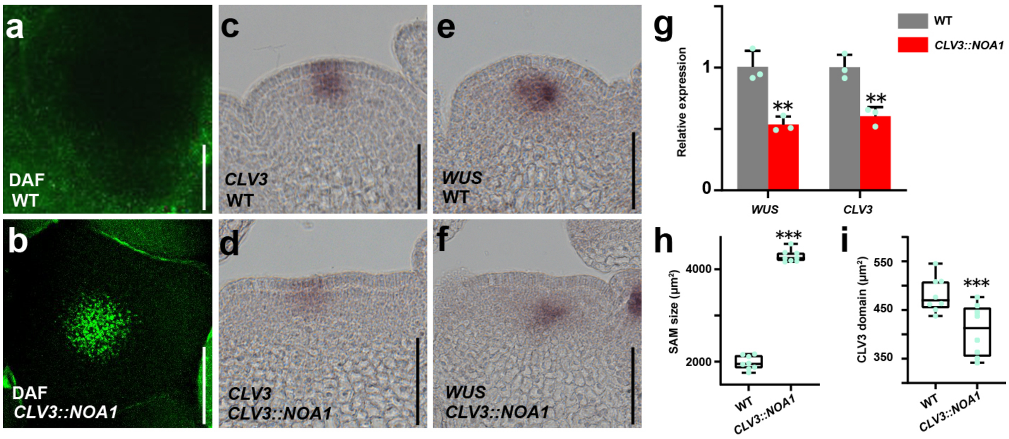


**Supplementary Fig. 6 | Nitric oxide promotes PZ fate.**

**a, b**, DAF-2DA staining in wild type (**a**) and *pCLV3::NOA1* transgenic plants (**b**). Scale bars, 50 µm. **c, d**, *CLV3* expression patterns in the wild type (**c**) and *pCLV3::NOA1* transgenic plants (**d**). **e, f**, *WUS* expression patterns in the wild type (**e**) and *pCLV3::NOA1* transgenic plants (**f**). **g**, RT-qPCR was performed to detect the expression levels of *WUS* and *CLV3* in the wild type and *pCLV3::NOA1* transgenic plants. The data are shown as mean ± s.d.; n = 3 biological replicates, two-tailed Student's t-tests, $^{**}P < 0.01$. All experiments were repeated at least two times (each time using multiple shoot apices from multiple plants), with similar results. **h, i**, Quantification the SAM size (**h**) and the size of stem cell region (**i**) in the wild type and *pCLV3::NOA1* plants. n = 8 independent plants, two-tailed Student's t-tests, $^{***}p < 0.001$. Scale bars, 50 µm.

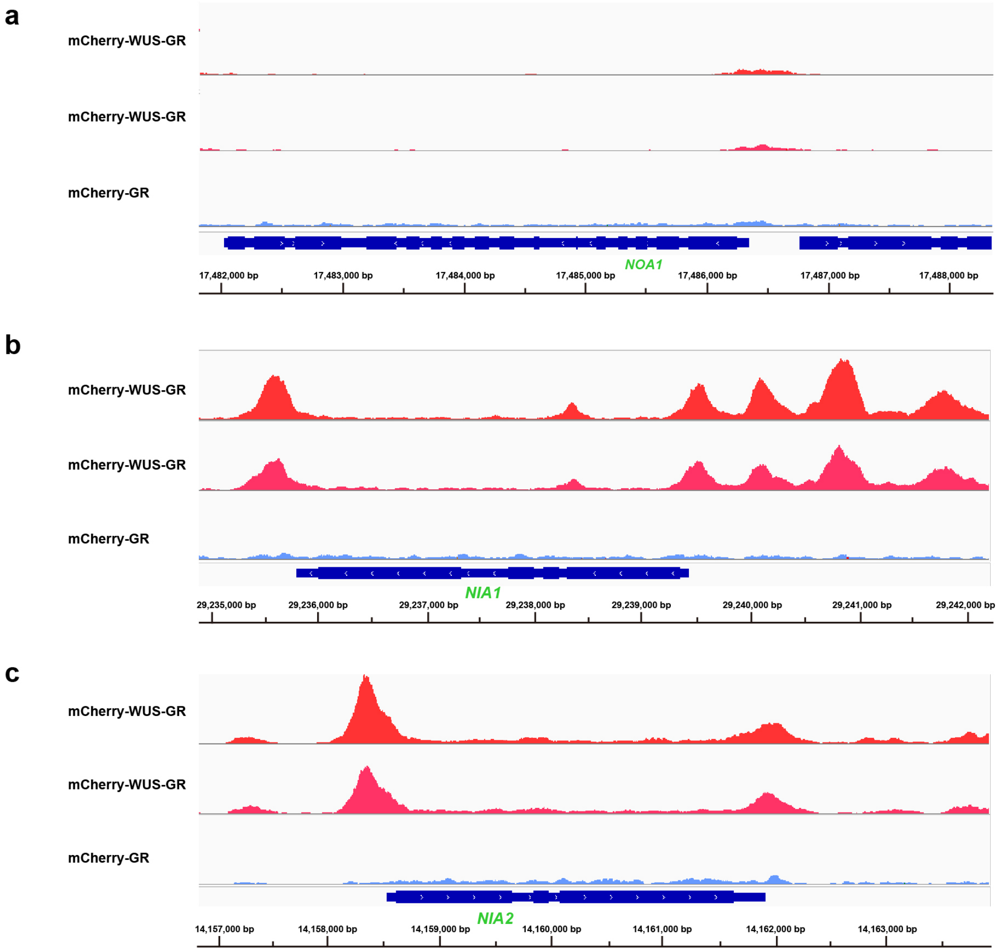


**Supplementary Fig. 7 | WUS associates with the promoters of NO biosynthesis genes.**

**a**-**c**, WUS Chip-seq traces of *NOA1* (**a**), *NIA1* (**b**) and *NIA2* (**c**) genomic regions after eChIP [2] on *pUBQ10::mCherry-GR-WUS* seedlings after Dex induction for 4 h.

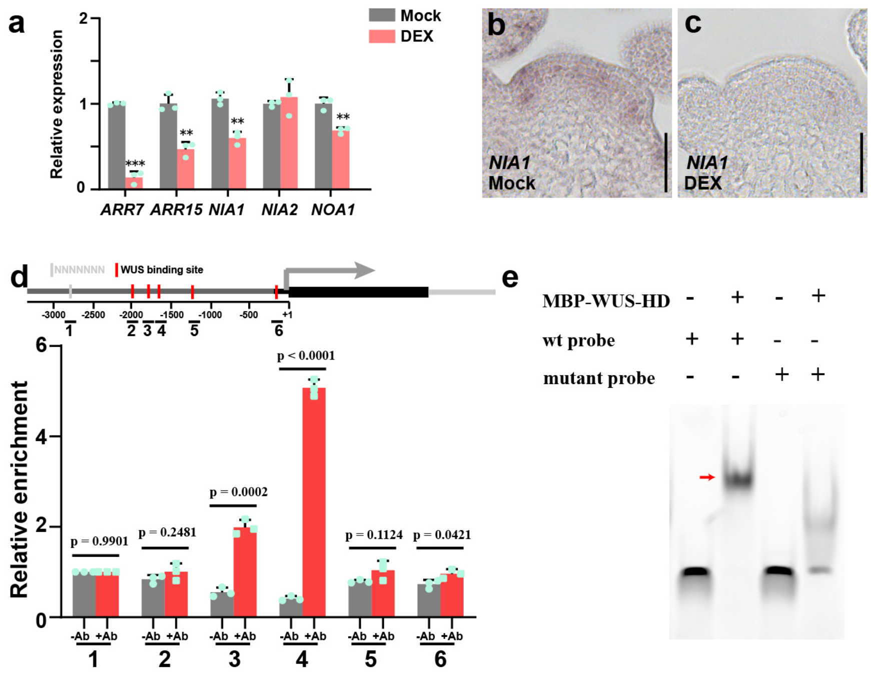


**Supplementary Fig. 8 | *WUS* directly represses NIA1 expression.**

**a**, Measurement of *ARR7*, *ARR15*, *NIA1*, *NIA2* and *NOA1* expression levels in the *pUBQ10::mCherry-GR-WUS* transgenic plants with and without Dex induction using RT–qPCR. The data are shown as mean ± s.d.; n = 3 biological replicates, two-tailed Student's t-tests, **p < 0.01, ***p<0.001. **b, c**, *NIA1* expression patterns in the *pUBQ10::mCherry-GR-WUS* transgenic plants with (**c**) and without (**b**) Dex induction. Scale bars, 50 µm. **d**, Enrichment of *NIA1* fragments after ChIP. Regions 1 to 6 are located in the *NIA1* promoter. –Ab, no antibody control; +Ab, with GR antibody. Red boxes indicates WUS binding motifs. The data are shown as mean ± s.d.; three independent experiments each for –Ab and +Ab. **e**, EMSA showing that WUS binds to the promoter (Region 4) of *NIA1 in vitro*. The red arrow indicates specific interactions. All experiments were repeated at least two times, with similar results.

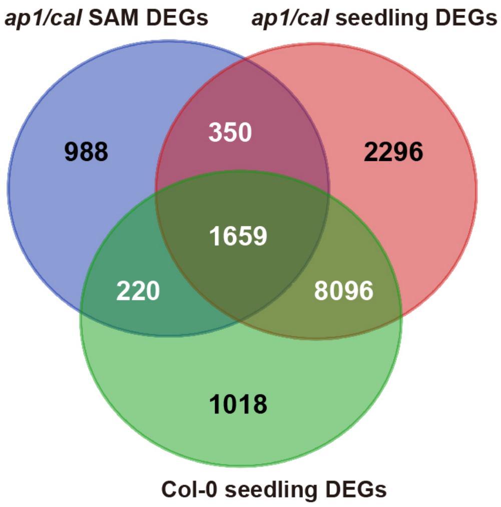


**Supplementary Fig. 9 | Comparison of transcriptional responses to NO in seedlings and SAMs of wild type and *ap1/cal* mutants .**

Venn diagram showing the overlap of DEGs in *ap1/cal* SAM and in Col-0, *ap1/cal* mutant seedling under 1mM SNP treatments.

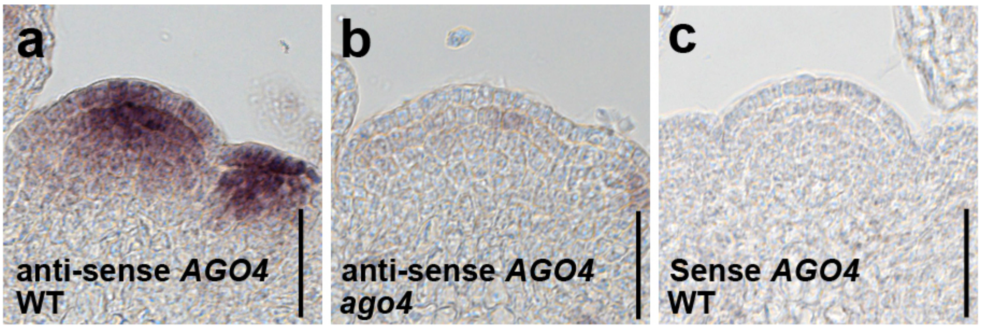


**Supplementary Fig. 10 | *AGO4* expression patterns in the SAM.**

**A**,**b**, *AGO4* expression patterns in wild type (**a**) and *ago4* mutant (**b**). **c**, Negative control using a *AGO4* sense probe. Scale bars, 50 µm.

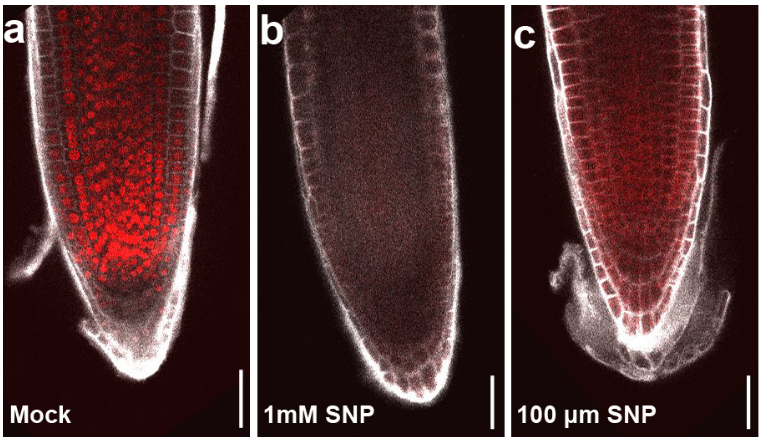


**Supplementary Fig. 11 | Nitric oxide affects AGO4 protein accumulation in the root apical meristem (RAM).**

**a**-**c**, AGO4-mCherry protein distribution in the RAM of *pAGO4::AGO4-mCherry/ago4* rescue plants after Mock (**a**), 1mM SNP (**b**) and 100 µM SNP (**c**) treatments. Scale bars, 50 µm.

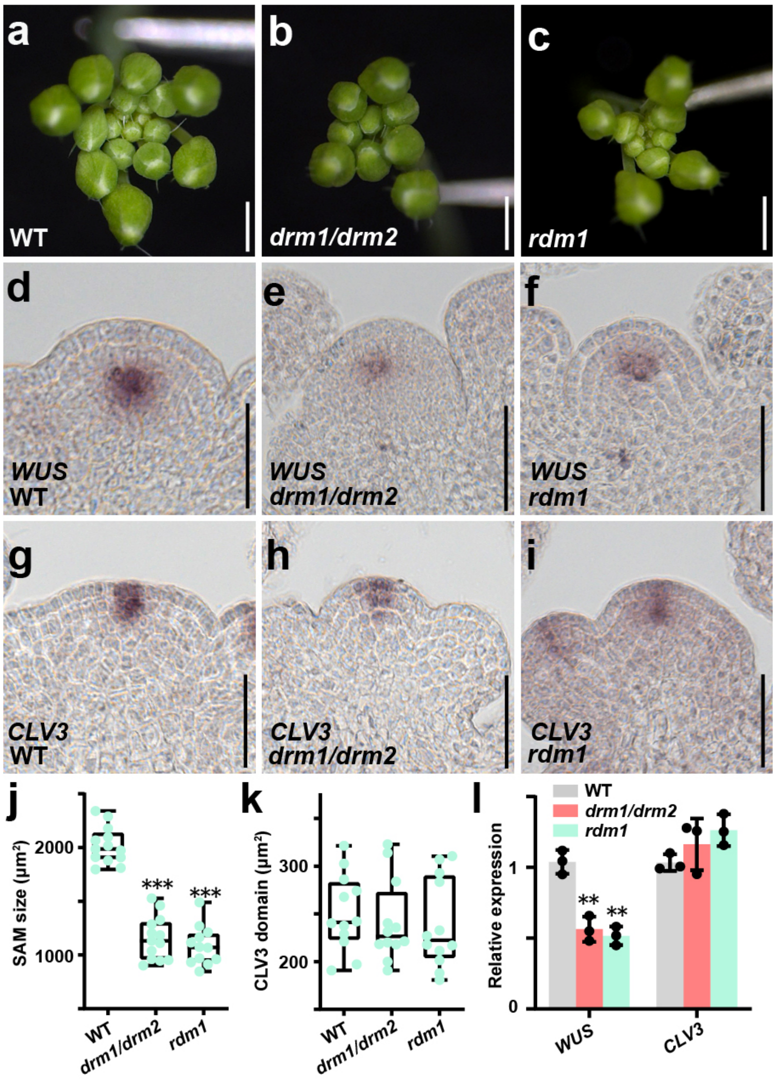


**Supplementary Fig. 12 | The RdDM pathway is required for stem cell homeostasis.**

**a-c**, Representative inflorescence phenotypes of wild type (**a**), *drm1/drm2* (**b**), and *rdm1* mutants (**c**). Scale bars, 1 mm. **d-f**, *WUS* expression patterns in wild type (**d**), *drm1/drm2* (**e**), and *rdm1* mutants (**f**). Scale bars, 50 μm. **g-i**, *CLV3* expression patterns in wild type (**g**), *drm1/drm2* (**h**), and *rdm1* mutants (**i**). Scale bars, 50 μm. **j, k**, Quantification of SAM size (**j**) and the size of stem cell region (**k**) in wild type, wild type, *drm1/drm2*, and *rdm1* mutants. n = 15, two-tailed Student's t-tests, ***$p<0.001$. **l**, RT-qPCR quantification of *WUS* and *CLV3* expression levels in wild type, *drm1/drm2* and *rdm1* mutants shown as mean ± s.d.; n = 3 biological replicates, two-tailed Student's t-tests, **$P<0.01$. All experiments were repeated independently at least two times on pools of apices of at least 30 plants.

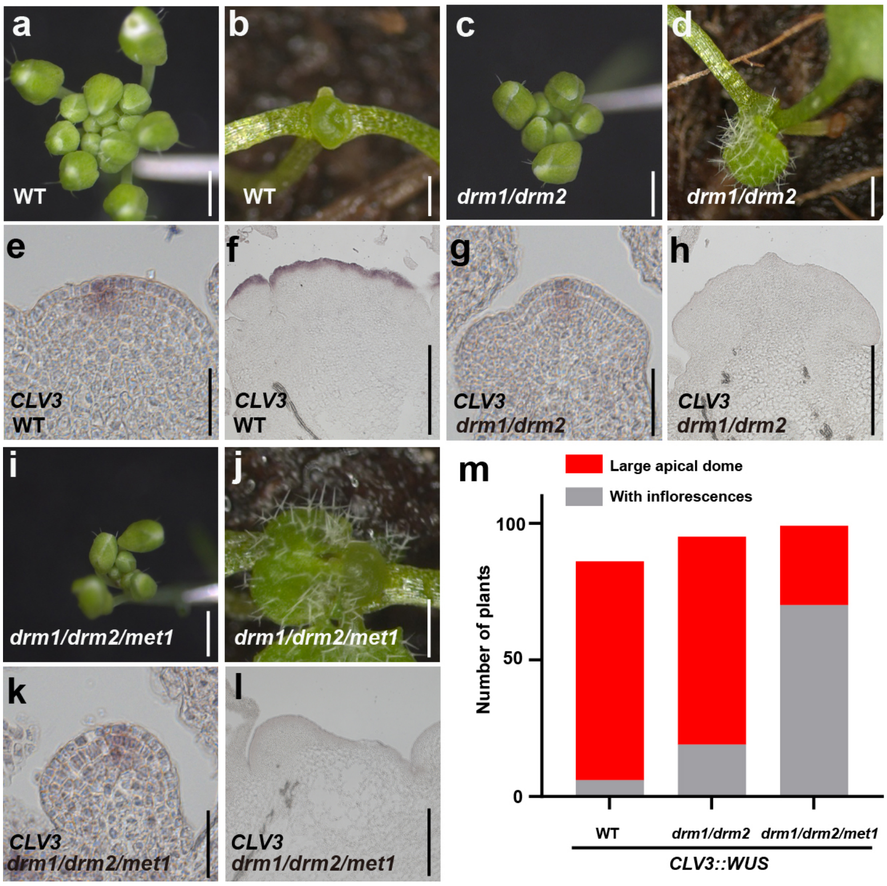


**Supplementary Fig. 13 | RdDM pathway is required for WUS protein function.**

**a-d**, Phenotypic analysis independent transgenic plants (T1) of *pCLV3::WUS* in wild type (**a, b**) and *drm1/drm2* (**c, d**) background showing two classes of phenotypes with (**a, c**) and without (**b, d**) inflorescences. **e-h,** *CLV3* expression patterns in the SAM of *pCLV3::WUS* in wild type (**e, f**) and *drm1/drm2* (**g, h**) background with (**e, g**) and without (**f, h**) inflorescences. **i, j**, Phenotypic analysis of independent T1 plants of *pCLV3::WUS* in the *drm1/drm2/met1* (**i, j**) triple mutants showing two classes of phenotypes with (**i**) and without (**j**) inflorescences. **k, l,** *CLV3* expression patterns in the SAM of *pCLV3::WUS* in the *drm1/drm2/met1* triple mutants with (**k**) and without (**l**) inflorescences. **m**, Phenotypic quantification of *pCLV3::WUS* in the wild type, *drm1/drm2* and *drm1/drm2/met1* background.

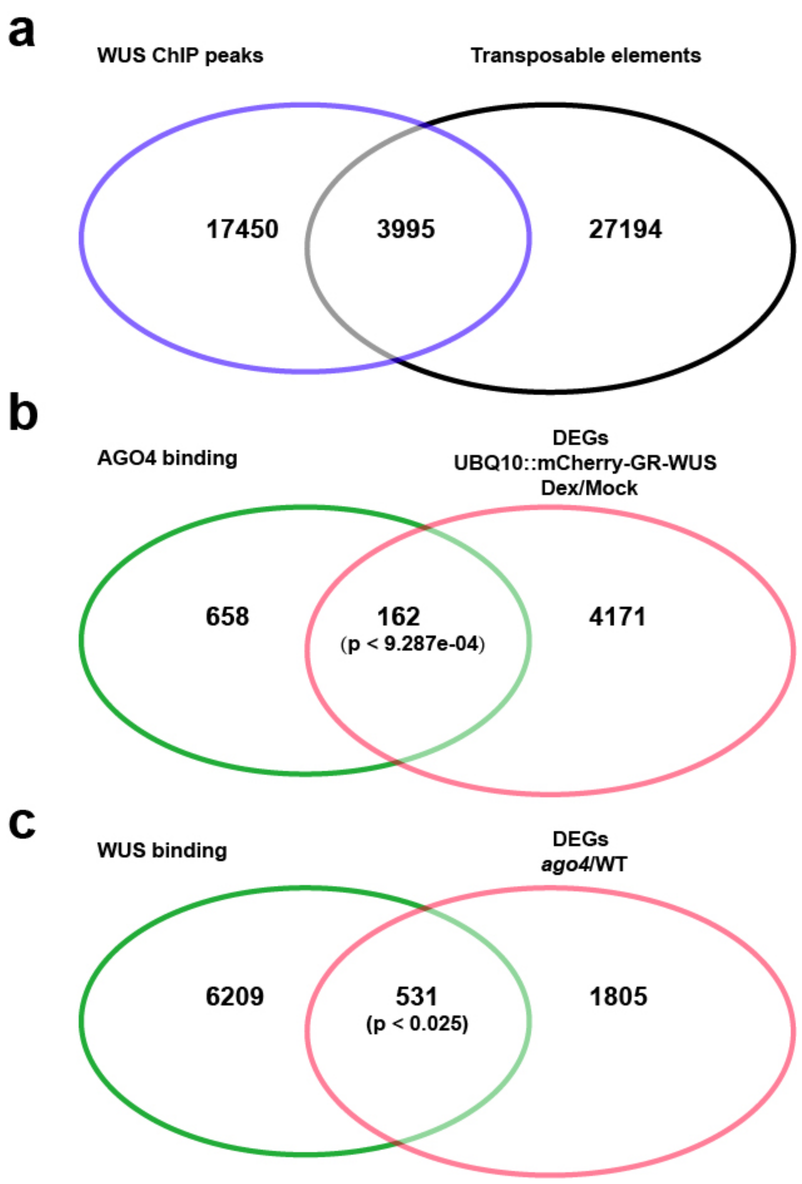


**Supplementary Fig. 14 | WUS regulates transposable elements.**

**a**, Venn diagram showing the overlap of annotated transposable elements and WUS binding peaks [4]. **b**, Venn diagram showing the overlap of AGO4 targets [5] and DEGs in response to WUS induction. **c**, Venn diagram showing the overlap of WUS targets and DEGs in *ago4* mutant vs wild type (GSE18928).

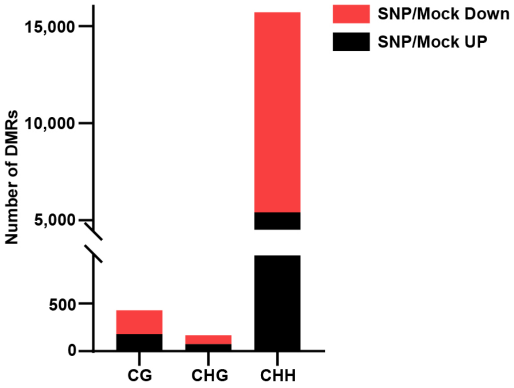


**Supplementary Fig. 15 | NO affects DNA methylation patterns in the SAM.**

Stratification of differentially methylated regions (DMRs) using our methylation data into CG (differentially methylated only in the CG context), CHG (differentially methylated only in the CHG context), and CHH (differentially methylated only in the CHH context) types.

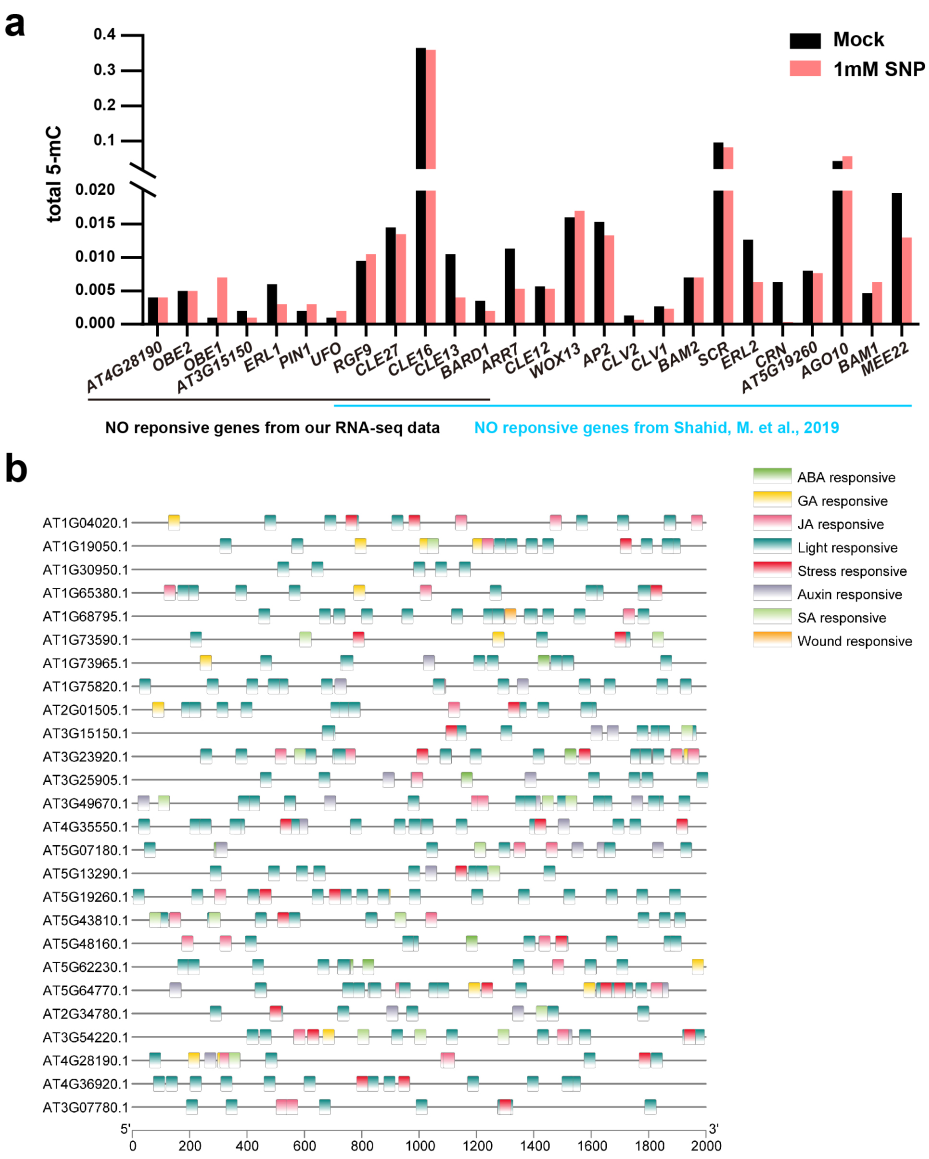

### Supplementary Fig. 16 | NO affects DNA methylation of stem cell related genes.

**a**, Total 5-mC DNA methylation levels at loci harboring NO responsive stem cell related genes in our dataset. **b**, Common responsive elements analysis of the NO responsive stem cell related genes.

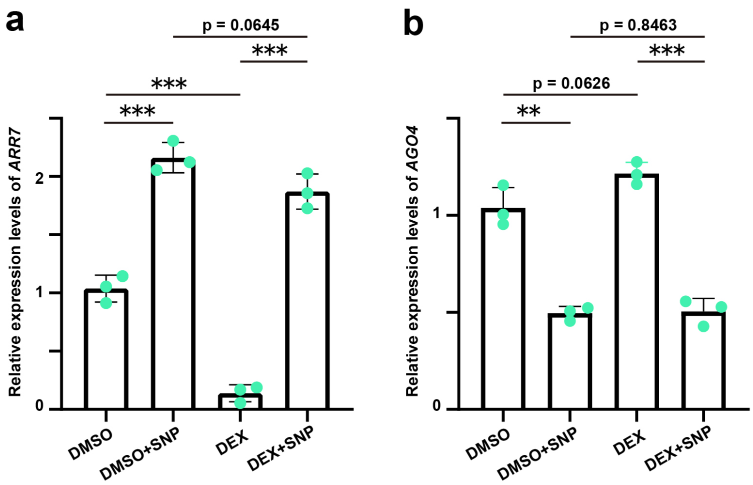


### Supplementary Fig. 17 | *AGO4* mediates NO signals to induce *ARR7* expression.

**a-b,** RT-qPCR measurements of *ARR7* (**a**) and *AGO4* (**b**) expression in *pUBQ10::mCherry-GR-WUS* plants with and without SNP/Dex induction. Data are shown as mean ± s.d.; n = 3 biological replicates, two-tailed Student's t-tests, **p < 0.01, ***p < 0.001.

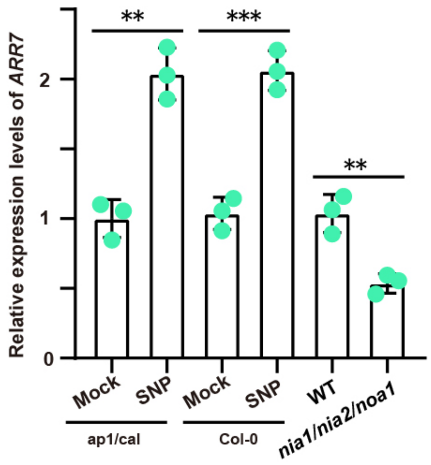


**Supplementary Fig. 18 | NO signaling induces *ARR7* expression.**

RT-qPCR measurements of *ARR7* expression levels in *ap1/cal* and Col-0 plants under SNP treatment, and *ARR7* expression levels in wild type and *nia1/2noa1* mutant. Data are shown as mean ± s.d.; n = 3 biological replicates, two-tailed Student's t-tests, $^{**}p<0.01$, $^{***}p < 0.001$.

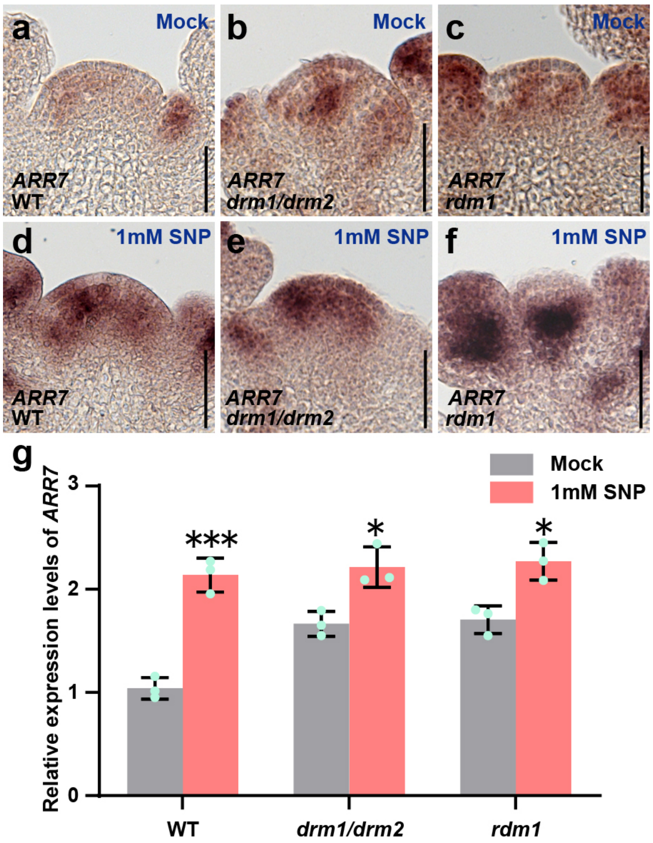


**Supplementary Fig. 19 | The RdDM pathway mediates NO signaling for proper expression of stem cell regulators.**

**a-c**, *ARR7* expression patterns in wild type (**a**), *drm1/drm2* (**b**) and *rdm1* (**c**) after mock treatment. **d-f**, *ARR7* expression patterns in wild type (**d**), *drm1/drm2* (**e**) and *rdm1* (**f**) after 1mM SNP treatment. **g**, Quantification of *ARR7* transcripts levels in wild type, *drm1/drm2* and *rdm1* after mock and SNP treatments shown as mean ±s.d.; n = 3 biological replicates, two-tailed Student's t-tests, *p<0.05, ***p<0.001. All experiments were repeated independently at least two times on pools of apices of at least 30 plants. Scale bars, 50 µm.

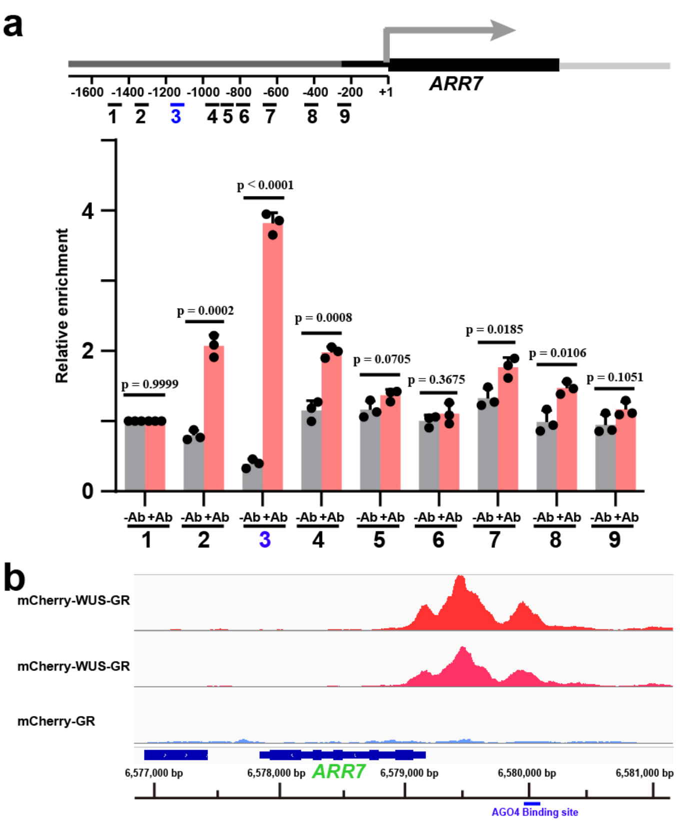


**Supplementary Fig. 20 | AGO4 binds *ARR7* promoter *in vivo.***

**a**, Enrichment of fragments of the *ARR7* regulatory region after ChIP using mCherry-AGO4 rescue plants. Regions 1 to 9 are located in the *ARR7* promoter. –Ab, no antibody control; +Ab, with mCherry antibody. The red box indicates WUS binding region. The data are shown as mean ± s.d.; three independent experiments we carried out each for –Ab and +Ab. **b**, WUS Chip-seq traces of the *ARR7* genomic region after eChIP [2] on *pUBQ10::mCherry-GR-WUS* seedlings after Dex induction for 4 h.

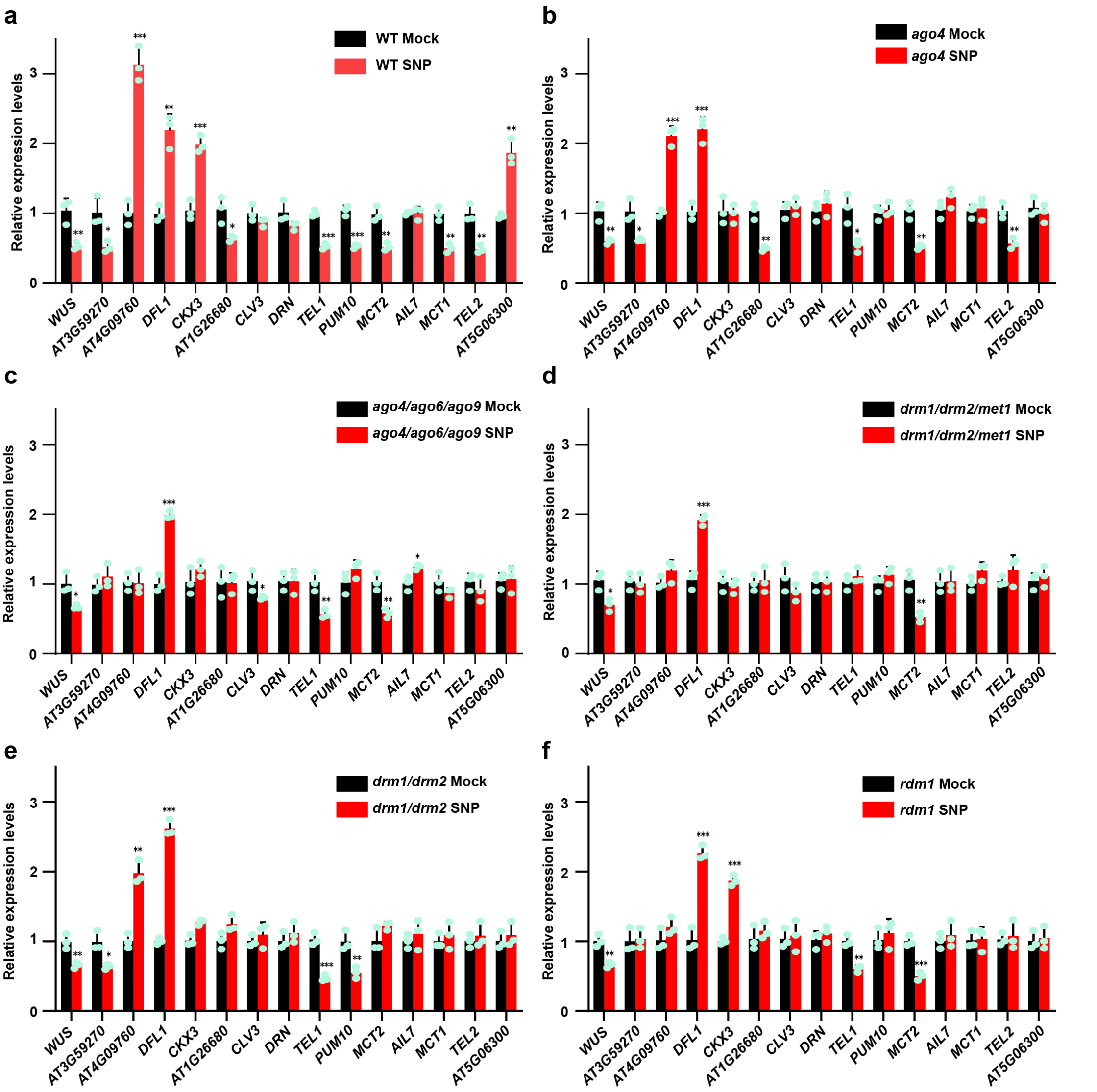


**Supplementary Fig. 21 | RdDM pathway mediates NO signaling in the SAM.**

**a-f**, RT-qPCR measurements of the expression levels of stem cell- and niche-specific genes in wild type (**a**) and in *ago4* (**b**), *ago4/ago6/ago9* (**c**), *drm1/drm2/met1* (**d**), *drm1/drm2* (**e**) and *rdm1* (**f**) mutants with and without SNP treatments. The data are shown as mean ± s.d.; n = 3 biological replicates, two-tailed Student's t-tests, *p<0.05, **p < 0.01, ***p<0.001.
